# Empirical study on software and process quality in bioinformatics tools

**DOI:** 10.1101/2022.03.10.483804

**Authors:** Katalin Ferenc, Konrad Otto, Francisco Gomes de Oliveira Neto, Marcela Dávila López, Jennifer Horkoff, Alexander Schliep

## Abstract

Software quality in computational tools impacts research output in a variety of scientific disciplines. Biology is one of these fields, especially for High Throughput Sequencing (HTS) data, such tools play an important role. This study therefore characterises the overall quality of a selection of tools which are frequently part of HTS pipelines, as well as analyses the maintainability and process quality of a selection of HTS alignment tools. Our findings highlight the most pressing issues, and point to software engineering best practices developed for the improvement of maintenance and process quality. To help future research, we share the tooling for the static code analysis with SonarCloud which we used to collect data on the maintainability of different alignment tools. The results of the analysis show that the maintainability level is generally high but trends towards increasing technical debt over time. We also observed that the development activities on alignment tools are generally driven by very few developers and are not utilising modern tooling to their advantage. Based on these observations, we recommend actions to improve both maintainability and process quality in open source alignment tools. Those actions include improvements in tooling like the use of linters as well as better documentation of architecture and features. We encourage developers to use these tools in order to ease future maintenance efforts, increase user experience, support reproducibility, and ultimately increase the quality of research through increasing the quality of research software tools.

## Introduction

Biology and medicine have seen a data-driven transformation with the advent of high-throughput experimentation, in particular high-throughput sequencing with ten thousands of sequenced genomes. Computational tools integrate advanced algorithms, data structures, and state-of-the-art methods from statistics and machine learning. However, the developers and users of research tools are generally experts in their respective fields and have little background in software engineering [1]. This hinders and delays the widespread adaptation of software development best practices, such as the usage of containers for deployment and reusability, within the bioinformatics community.

High Throughput Sequencing (HTS) is a field of bioinformatics, which includes many of the state of the art approaches for analyzing DNA or RNA sequences generated by sequencing instruments [2]. The analysis of HTS relies heavily on software tools developed by various research groups. Hundreds of HTS pipelines are built and employed in laboratories around the world to analyse the results of HTS applications and draw conclusions in a wide variety of fields of biology. The applications include measuring biodiversity, developing novel medical treatments, or diagnosing cancer subtypes in patients. Despite their widespread usage and significance, the quality of these bioinformatics tools has been poorly investigated from the software engineering point of view, with some notable exceptions.

Previous findings indicate that many software qualities were not prioritised during the development process of many of these tools. For example, the testability of these tools is not adequately addressed [3], the accessibility and installability of bioinformatics software was found poor [4] even though tools like Bioconda have recently improved the situation. Furthermore, user inputs and dependencies are not checked properly, and status logs are missing [5, 6]. These shortcomings result in end users unsatisfied with the user experience of these tools [7]. Although users can choose their favourite from a multitude of software tools available to solve each step in a standard pipeline, the compatibility of these tools is not always guaranteed [8]. Furthermore, the creation and usage of pipelines often requires the knowledge of at least one of several pipeline management frameworks [9]. As a result, experimental researchers often rely on bioinformaticians to create and maintain their pipelines, or even restrict themselves to problems they feel comfortable solving with the tools they are already familiar with [7].

Most of the issues bioinformatics software development is facing are known and studied in the field of software engineering. Knowledge exchange and guidelines that are rooted in the best practices of software engineers can prove to be useful to practicing bioinformaticians. However, to achieve transferability of software engineering practices to scientific software development, issues within scientific communities need to be correctly identified and field-specific constraints should be considered. Killcoyne and Boyle claim that the nature of the *ad hoc* scientific approach and the structured software engineering approach is conflicting [10]. We argue that several tools used in the bioinformatics community are in fact in the maintenance phase of their life cycle, where the *ad hoc* solutions should be replaced with a robust code. The maintainability of scientific software has a direct impact on the reproducibility of research findings, as well as on the time required for the developers to fix bugs and consequently, on the waiting time for the user to run a updated tool when encountering a breaking issue.

Therefore, we aimed to characterise some of the well-known bioinformatics software tools, which are in the maintenance phase of their life. First, we investigated whether a selection of currently available, established, widely used, open source bioinformatics tools address previously identified software quality issues including maintainability, usability, and documentation. To this end, we evaluated the status of 20 HTS software tools across 7 categories to get a broad overview of their quality in light of previous research findings. Next, we investigated the code-level artefacts and software development practices of 13 different, but comparable tools focusing on maintainability. In particular, we selected alignment tools which are indispensable parts of any HTS workflow. These tools are responsible for identifying the location of short DNA fragments (reads) to the reference genome [11]. This process is called alignment or mapping, and the tools are the aligners or mappers. Finally, we analysed the evolution of 3 open source aligners to identify trends in maintenance and code quality.

Our aim was to identify key issues in widely used bioinformatics software tools from the software engineering perspective, especially to evaluate to what extent software engineering best practices are followed during software maintenance. We acknowledge the lack of resources available for software maintenance, however, we would like to stress the importance of good quality software for good quality research. Our experimental findings support the necessity of following the guidelines found in literature [8, 12–15]. This study is organised into three parts, one for each research question (RQ).

**RQ1:** What are the software quality characteristics of tools used in workflow pipelines in HTS data analysis?
**RQ2:** What is the state of software and process quality in popular open source alignment tools?

**RQ2.1:** How maintainable is the source code of alignment tools?
**RQ2.2:** What characterises the development activity of open source projects for alignment tools?
**RQ3:** How does the maintainability of alignment tools evolve? Are there trends in the software quality between the versions of a selected alignment tool?

## Methods

### Characterization of HTS tools

The data collection was performed at the Bioinformatics Core Facility, Sahlgrenska Academy at the University of Gothenburg. The Core Facility employs bioinformaticians and statisticians who support a wide variety of research projects and organize workshops and training sessions. To this end, 20 standard, widely used tools (Table 1) were selected from the HTS pipelines used in 2019 at the Core Facility. These tools are fully developed and in the maintenance phase of their life cycle, thus their characterization can shed light on the status of tools used by biological data analysts.

**Table 1.**
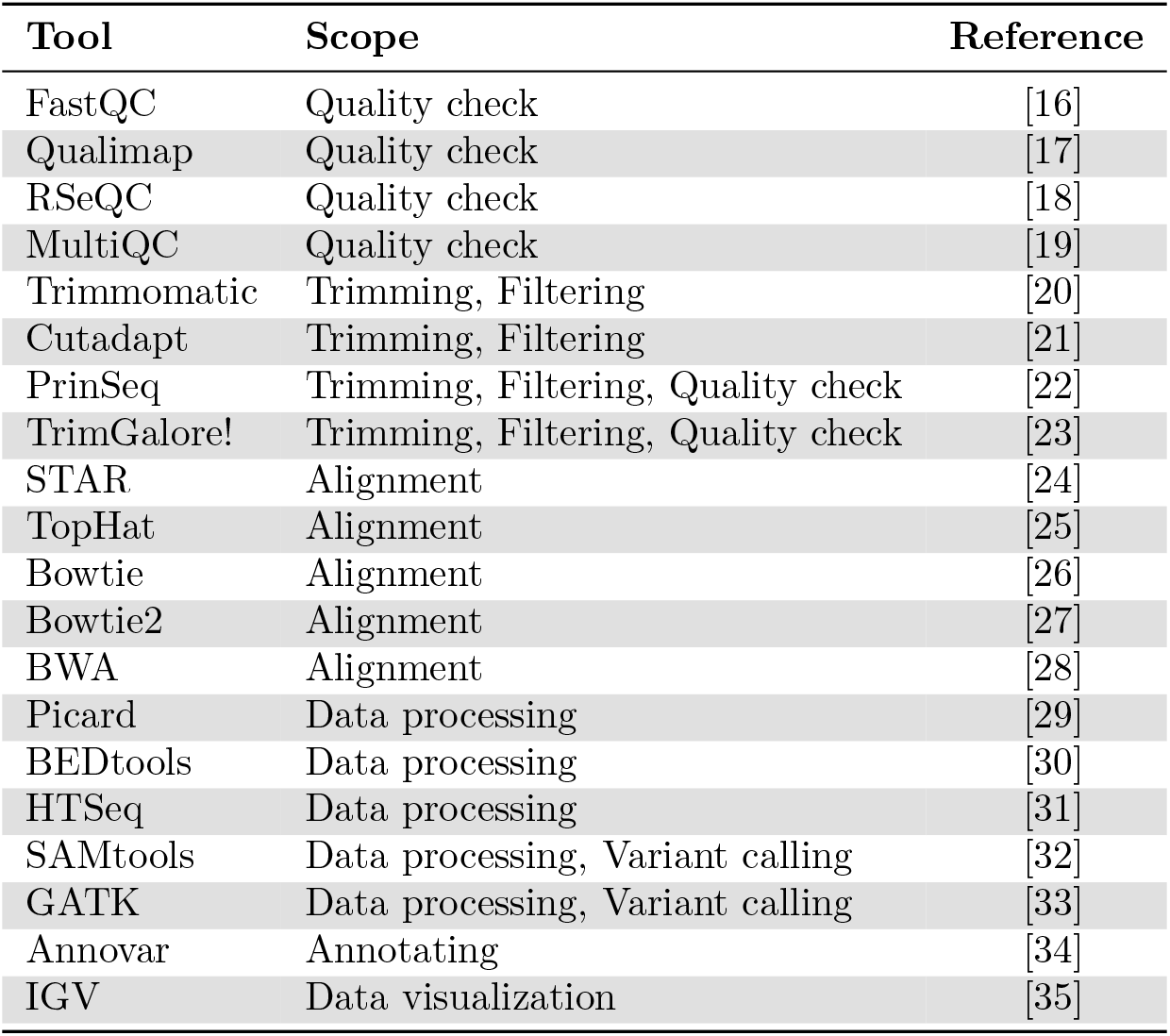
List of Tools Selected for RQ1 with the Scope of Characterisation Applied to Each.

The main data source for RQ1 (characterization of tools) was the available documentation, manual and GitHub repositories of the selected tools. Additionally, a test installation and one or several test runs were performed using publicly available DNA-seq (SRA: SRX3122951) and RNA-seq (SRA: SRP130955) data, using Ubuntu 16.04 version. The software tools were characterized using 7 criteria selected from literature [4, 7–9, 14, 36–38] and focusing on areas important to the everyday work of users of these tools. The criteria are: 1) integration into the pipeline, 2) maintenance, 3) support, 4) usability, 5) documentation, 6) installability, and 7) dependency management. For each criterion, a 3-level quality scale was defined (Table 2) and applied following tests and review of the documentation.

**Table 2.** Definitions of the different levels for the 7 characteristics of HTS tools used in this analysis.

| PIPELINE INTEGRATION |  |
| --- | --- |
| 1 | can run directly using any input and output files of standard format |
| 2 | requires specific input file name (e.g. *_1.fq and *_2.fq) |
| 3 | requires non standard file format or non standard changes within the input file |
| MAINTENANCE |  |
| 1 | latest commit no more than 1 year before this analysis (2020) |
| 2 | more than 1, but less than 5 years inactivity before this analysis |
| 3 | more than 5 years inactivity |
| SUPPORT |  |
| 1 | has its own active GitHub issue / forum / bug report page |
| 2 | documentation about frequent errors and / or email support, but inactive issue page |
| 3 | user needs to rely on external forums or resources |
| USABILITY |  |
| 1 | has a graphical user interface in Galaxy or standalone, test finished |
| 2 | only command line interface AND running requires 1 command for a single task, OR test crashed with useful error messages |
| 3 | only command line interface AND running requires at least 2 commands for a single task, OR test crashed with hard to understand or missing error messages |
| DOCUMENTATION |  |
| 1 | sufficient and necessary information for usage, easy to navigate |
| 2 | short, not well organized or hard to read |
| 3 | too short for effective usage, requires additional external resources |
| INSTALLATION & DEPENDENCIES |  |
| 1 | self-contained, i.e. sufficient information on the website and/or in the documentation |
| 2 | requires admin rights or external help |
| 3 | requires external tools |

### Data collection for RQ2 and RQ3

The most widely used sequencing technologies produce short (30-100 base pairs long) DNA or RNA sequences (reads) which are then aligned (mapped) to the reference genome of the species of interest. The first step in a standard analysis pipeline is the quality check of the sequenced reads, which can be followed by a quality-based filtering or trimming. Next, the reads are mapped to the reference genome. The mapping quality is investigated by the bioinformatician before proceeding to subsequent, application-specific data analysis. The analysis process relies on multiple tools and frequent quality checks which drives the subsequent steps, such as the abortion of the protocol or adjustment in the parameters of the tools.

For RQ2, we decided to focus on a single step in the investigated HTS pipelines. We chose the mapping step, as several mappers have been developed and utilized in the majority of HTS applications. Most mappers are written in the `C` or `C++` languages allowing for fair code-level comparison. We aimed to include several widely used open source mappers [39]. This selection resulted in a total of 13 different mappers, all with their code available on GitHub, being analysed. Table 3 lists the selected mappers and provides a link to the each project’s source code.

**Table 3.**
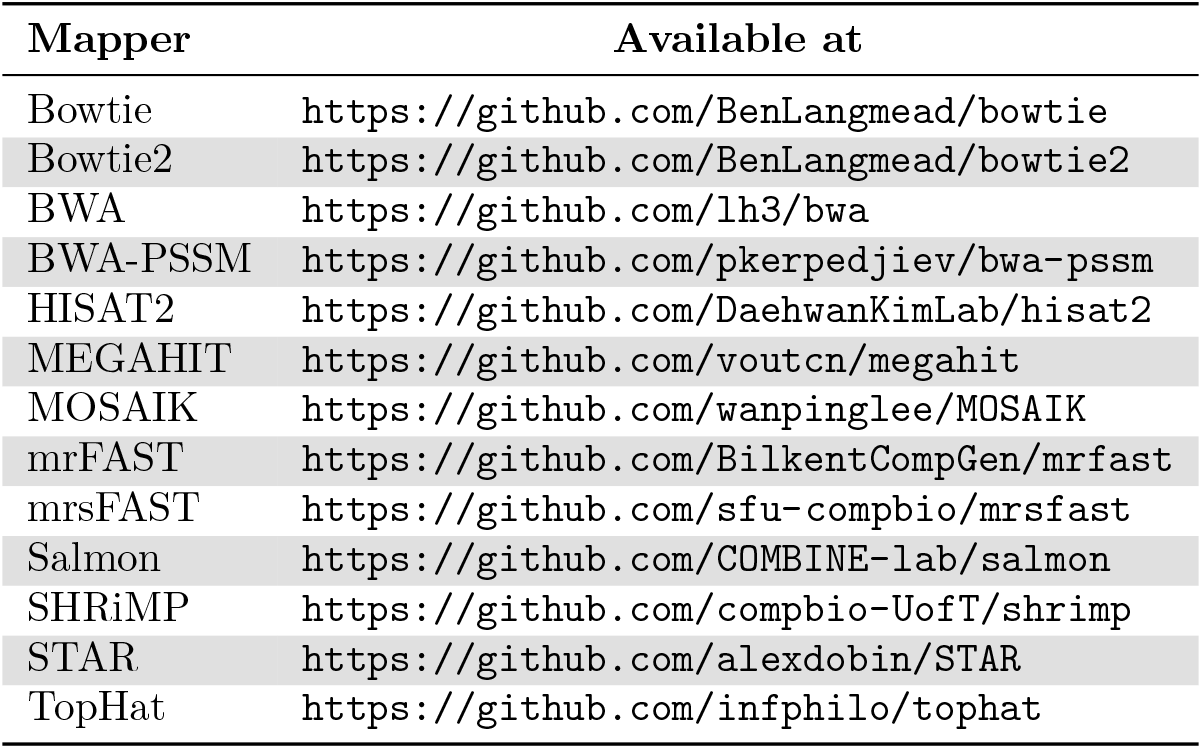
List of Mappers Selected for Analysis.

| Mapper | Available at |
| --- | --- |
| Bowtie | https://github.com/BenLangmead/bowtie |
| Bowtie2 | https://github.com/BenLangmead/bowtie2 |
| BWA | https://github.com/lh3/bwa |
| BWA-PSSM | https://github.com/pkerpedjiev/bwa-pssm |
| HISAT2 | https://github.com/DaehwanKimLab/hisat2 |
| MEGAHIT | https://github.com/voutcn/megahit |
| MOSAIK | https://github.com/wanpinglee/MOSAIK |
| mrFAST | https://github.com/BilkentCompGen/mrfast |
| mrsFAST | https://github.com/sfu-compbio/mrsfast |
| Salmon | https://github.com/COMBINE-lab/salmon |
| SHRIMP | https://github.com/compbio-UofT/shrimp |
| STAR | https://github.com/alexdobin/STAR |
| TopHat | https://github.com/infphilo/tophat |

### Static code analysis

*Maintainability* is one of the software qualities listed in ISO/IEC 25010 [40]. In the model it is constructed from *modularity*, *reusability*, *analysability*, *modifiability*, and *testability* [40]. According to Riaz *et al*., it can be summarised as “the ease with which a software system can be modified” [41]. Maintainability is an indicator for the amount of work needed to understand, reuse, and refactor a software component or project [42]. Making changes or fixing bugs takes less effort in a project with good maintainability [42]. Maintainability is directly affected by the quality of the applied development process. Sing and Gautam describe the connection between the two, especially how the four development activities (i.e., requirements, design, coding, and testing [43]) included in any type of development process (e.g. waterfall, scrum) can impact the maintainability of the end product [44]. Table 4 shows exactly what requirement characteristics, design attributes, coding factors, and testing parameters can be utilised to improve maintainability.

**Table 4.**
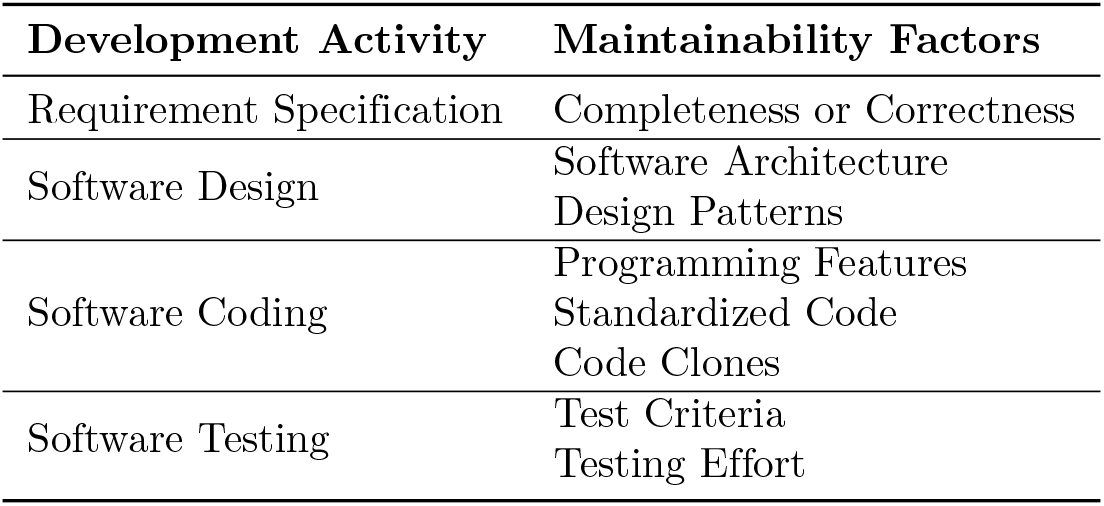
Factors that can be used to improve maintainability during software development activities (adapted from [44])

| Development Activity | Maintainability Factors |
| --- | --- |
| Requirement Specification | Completeness or Correctness |
| Software Design | Software Architecture<br>Design Patterns |
| Software Coding | Programming Features<br>Standardized Code<br>Code Clones |
| Software Testing | Test Criteria<br>Testing Effort |

We used static code analysis to obtain the necessary measures for the maintainability evaluation of HTS mappers. According to Riaz *et al*. [41] there is a multitude of different metrics in the existing research which are used to quantify the maintainability of software projects. In this study, maintainability is inspected based on code smells and technical debt. Code smells describe the presence of bad programming and bad code design in a software artefact [45]. Code smells can cause maintainability issues, decrease comprehensibility, and cause concern to professional developers [46]. They can be detected with automated tools for static code analysis and can therefore easily be target for refactoring and improving code quality [45, 47]. The concept of technical debt is based on a metaphor comparing the future work that quality issues in production software cause to indebting yourself. The software can be released immediately without resolving the quality issues. Not having fixed the issues can however result in additional effort (potentially more than would have been necessary to fix the original issue) becoming necessary in the future [48, 49].

SonarQube is one of the most widely used solutions for static analysis of software quality [50–52]. For this work, we decided to use the online version of the tool SonarCloud^1^, as it is freely available, suitable for `C` and `C++` languages, supports continuous integration, and enables direct analysis of code stored in GitHub and Bitbucket repositories. These properties make SonarCloud appropriate for this research and for future usage within the bioinformatics community.

We used manual analysis of SonarCloud, which is straightforward and can easily be repeated on a multitude of different tools. To enable reproducibility, we decided to provide a Docker image for the analysis. All dependencies for the project build and the manual analysis are contained in the image and the commands for analysis are also provided. The Docker image for replicating the analysis is publicly available on GitHub (https://github.com/konradotto/sonar-analysis). This image has also been published on DockerHub (https://hub.docker.com/r/konradotto/sonar-analysis) and can therefore be fetched from there directly as konradotto/sonar-analysis.

As our analysis is focusing on maintainability, we selected the following measures provided by SonarCloud:

- Size: lines of code (LOC)
- Number of code smells
- Technical debt (estimated time to fix issues)
- Debt ratio (technical debt divided by the estimated time to create a debt-free solution from scratch)
- Maintainability rating
- Number of files
- Number of functions

The size of the various projects is important to allow a fair comparison of the absolute measures provided by SonarCloud. By normalizing for the LOC, it is possible to compare projects based on their inherent number of code smells and technical debt. The debt ratio and maintainability rating are directly related and already relative measures that take the project size into account. This maintainability rating allocates letter grades (*i.e*. in descending order: A, B, C, D, E) to projects based on their debt ratio. Finally, the number of files and the number of functions provide superficial, but valuable insight on the modularity and design-level differences between the projects.

### Development activity

To analyse the development activity on open-source mappers for HTS, artefacts from the online versioning systems used for the project have been consulted. The GitHub REST API has been used to collect those artefacts through the *Requests* Python HTTP library [53]. The contributors’ GitHub aliases were anonymised for ethical reasons. For subsequent analysis *ad hoc* Python scripts were used.

The measures about the contribution process that we are interested in for the analysis are the following:

- Number of major contributors (bus count)
- Total number of commits
- Temporal distribution of commits
- Number of releases or tags
- Commits per release or tag
- Open issues
- Ratio of open issues versus all
- Duration until issues are fixed

These measures have been chosen due to their perceived significance for the quality of the development process. Their importance is supported by the models that have been designed to predict “socio-technical issues” in [54] and [55]. The term “bus count” indicates how many developers have in-depth knowledge of a project, system or component (*e.g*. in a distributed system); it is the minimum number of developers that would have to suddenly disappear to endanger a smooth continuation. Tags are defined as snapshots in time. They are closely related to releases, and can be used to follow the development of the code throughout time. The high number of commits per tagged release for some of the other projects can be seen as a sign of missing direction in the development or underutilisation of releases. In either case this means that development and improvements are not sufficiently broken down into small increments (*i.e*. releases) that are communicated to the users.

## Results

### Characterisation of HTS tools

We summarised our findings on Table 5. The quality of the evaluated tools differed across categories. In line with the findings of Mangul et al. [4], the majority of the tools performed poorly on the scales of installability and dependency management, but most achieved good scores in several other categories.

**Table 5.**
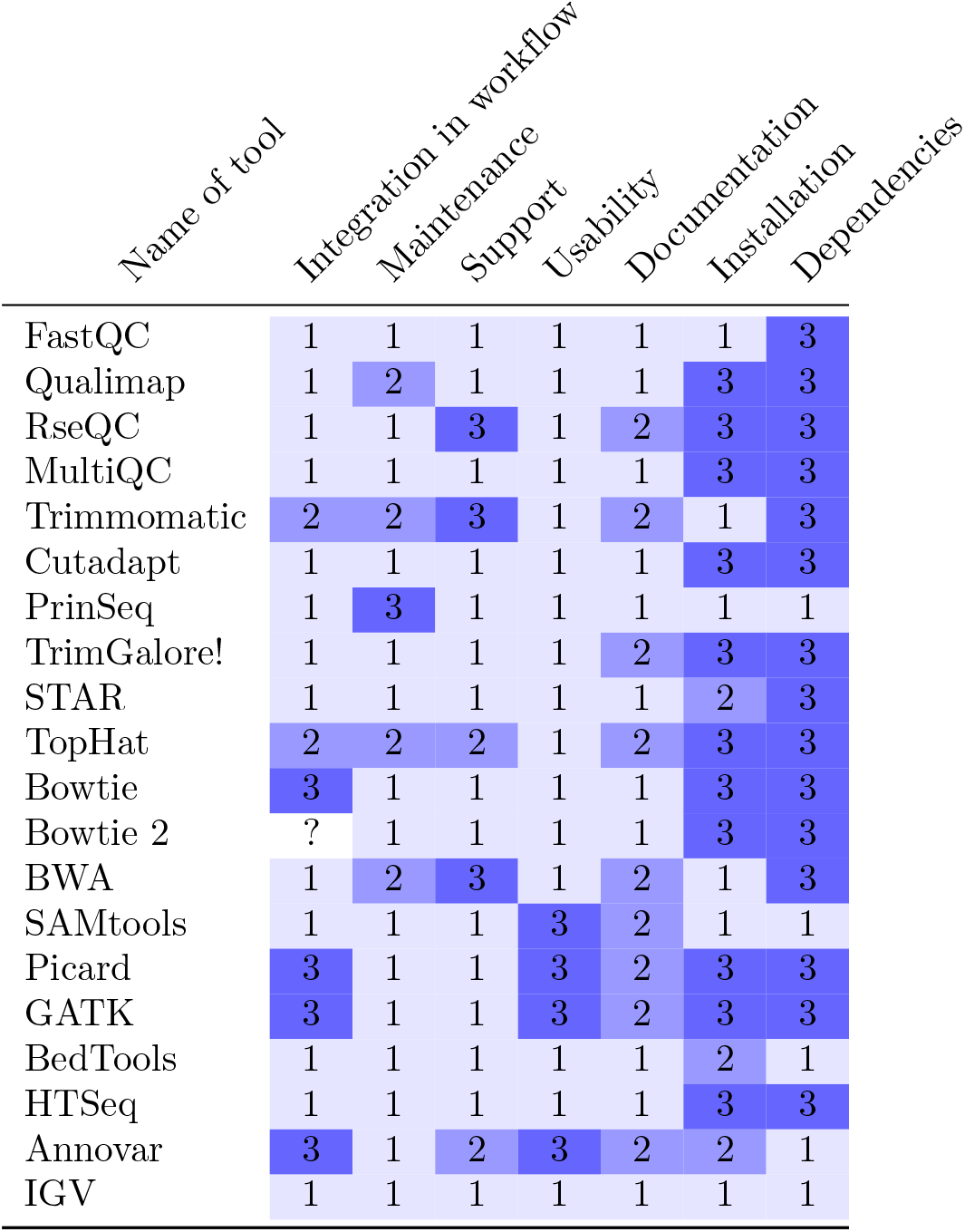
Scores of the analysed HTS tools on the scale defined in Table 2. 1: highest quality, 2: medium quality, 3: lowest quality.

Integration of tools into workflows relies on scripts written by the users. For example, running the same steps for several inputs, setting file structure and naming conventions, parallelisation, downloading database dependencies are rarely included in the tools. Workflow managers, such as Snakemake [56] or Nextflow [57] supports some of these tasks. However, we noted some incompatibilities between tools which require in-depth knowledge of them. One example is the presence (chrN) or absence (N) of chromosome prefixes in input/output files, i.e., compatibility with UCSC or Ensembl style reference genomes. Similarly, some tools are compatible with a specific subsequent tool, such as Trim Galore! being compatible with Bowtie1 by performing an additional base trimming from the input sequences [23]. This extra step is not necessary for the compatibility with other tools. We also noted that some steps can be omitted from the pipeline. For example, the BWA and Bowtie2 tools provide soft and hard clipping, thus removing the need to rely on the trimming of reads. However, in some applications this step is recommended.

Most of the investigated tools have been actively maintained at the time of the analysis (2020) with the exception of PrinSeq [22]. To investigate trends in the maintenance of tools, we performed an in-depth analysis of short sequence mappers in RQ3. Most tools are supported at their own website or other platforms visited by the creators. We noted that complete lack of support was rarely the case within the bioinformatics community. Platforms such as Biostars [58] and StackExchange [59] hosts extensive community knowledge for troubleshooting.

When considering usability, it is important to note that most tools are only available with command line interfaces, limiting their usage to bioinformaticians or scientists with knowledge in bash scripting. We argue that investing in a graphical user interface would increase the learnability of these tools and (potentially at the expense of a reduced number of available settings) would enable more researchers to utilize them in their work. One such example is the Galaxy tool that hosts a collection of independently developed tools and provides a graphical interface to them [60]. Indeed, it enables wider usage of the tools and independence from bioinformatics support.

Additionally, we found poor error management of several of the tools which further limits their usage to bioinformatician experts. For example, we tested Annovar with Variant Call Format (VCF) input format instead of the standard TXT format and it resulted in empty output file instead of an error message. Another example was Picard, which produced a 43 line long error log for a single error (“SAM file doesn’t have any read groups defined in the header.” - caused by incompatibility between Picard and SAMtools). In the same error message another error was mentioned (“StatusLogger Log4j2 could not find a logging implementation. Please add log4j-core to the class path”) which did not cause break in the code, but caused confusion when trying to find the main issue. A similar issue was observed when GATK was tested. The RealignerTargetCreator, IndelRealigner and BaseRecalibrator functions were run in sequence on the same input file, and the subsequent errors masked the original formatting error. Additionally, we noted that some tools have the potential to be merged into a single step. Especially in GATK functions such as VariantRecalibrator and ApplyRecalibration, or RealignerTargetCreator and IndelRealigner depend on each other in a linear order and thus can be merged into one function for the user.

We observed various types of documentation. We found documentations following the guidelines of Lee [12]. For example, the documentation of SAMtools includes common usage, and the documentation of GATK and BWA are written in an easy-to-read language. BEDtools features interesting usage examples from the community, which is a nice utilization of online forums and community resources. On the other hand, we also found that some documentation are not compliant to the guidelines. For example, Annovar’s documentation is unstructured, and it is hard to navigate in the documentation of Bowtie2. We noted that it is easier to find usage examples of Bowtie2 on forums than in the documentation.

The low mark (score 3) of dependency management was mainly due to their depencies on an external software which is not included in their release. For example, TrimGalore! is a wrapper around Cutadapt and FastQC, but installation does not include these two tools. This means that additional time and effort is required from the user to install dependencies separately from a third party website. In some cases, such as in TopHat, we found the requirement that another tool should be in the PATH, requiring additional steps.

In line with the finding of Mangul et al. [4], the installation of the investigated tools was longer than expected due to the lack of information or additional installation of dependencies from third party websites. Furthermore, several of the tools requires root privileges (e.g., STAR and GATK), which hinders the fast exploration and development of applications, as clusters or even personal working computers might be managed by IT personnel.

RQ1 We observed the following points about the characterization of tools:

- Most investigated tools have good documentation and are maintained at the time of analysis
- Several tools have score 3 (poor quality) on support, workflow integration, and usability. These shortcomings require additional time to spent and glue code to apply for the end user
- Most investigated tools have score 3 (poor quality) on installation and dependency management due to the need for external help or third party tools. We expect this issue to be minimized with the usage of Bioconda

### Software and Process Quality of Mappers

#### Software Maintainability

Table 6 summarises the results of the static code analysis for all mappers that we examined. Aside from *BWA-PSSM*, *mrFAST*, and *MOSAIK*, all repositories were analysed at the latest tagged version available at the time of analysis. We made exceptions for *BWA-PSSM* and *mrFAST* because the most recent tag was significantly older than the most recent changes by over 1 year and nearly 4 years respectively. *MOSAIK* had no tags at all. Therefore the latest commit was analysed for these three projects instead.

**Table 6.** Overview of the Maintainability and Size Metrics Collected from SonarCloud. The entries are sorted by the debt ratio of the projects in increasing order.

| Project | Version | Language | LOC | Files | Functions | Code Smells | Debt | Debt Ratio | Rating |
| --- | --- | --- | --- | --- | --- | --- | --- | --- | --- |
| Bismark | v0.16.2 | HTML | 4773 | 6 | 0 | 57 | 4 h 45 min | 0.2 % | A |
| MEGAHIT | v1.2.9 | C++ | 22 795 | 127 | 2176 | 1752 | 32 d | 2.3 % | A |
| STAR | 2.7.8a | C/C++ | 45 619 | 321 | 466 | 5231 | 67 d | 2.4 % | A |
| TopHat | v2.1.2 | C++ | 243 037 | 745 | 5619 | 39 618 | 424 d | 2.8 % | A |
| BWA-PSSM | latest | C | 15 732 | 73 | 538 | 1925 | 34 d | 3.6 % | A |
| HISAT2 | v2.2.1 | C | 113 373 | 230 | 3369 | 12 174 | 270 d | 3.8 % | A |
| Salmon | v1.4.0 | C++ | 21 638 | 47 | 446 | 3573 | 54 h | 4.1 % | A |
| BWA | v0.7.17 | C | 14 500 | 61 | 544 | 1937 | 37 d | 4.2 % | A |
| mrFAST | latest | C | 4960 | 15 | 94 | 620 | 13 d | 4.3 % | A |
| Bowtie2 | v2.4.2 | C++ | 63 384 | 147 | 2462 | 7514 | 168 d | 4.3 % | A |
| SHRiMP | v2.0.3 | C++ | 14 268 | 65 | 323 | 2408 | 42 d | 4.7 % | A |
| mrsFAST | v3.4.2 | C | 5831 | 20 | 130 | 992 | 18 d | 5.0 % | B |
| MOSAIK | latest | C++ | 30 924 | 172 | 989 | 5577 | 96 d | 5.0 % | B |
| Bowtie | v1.3.0 | C++ | 41 078 | 112 | 1858 | 7798 | 142 d | 5.5 % | B |
| Median<sup>†</sup>: |  |  | 22 795 | 3.4<sup>§</sup> | 29.7<sup>§</sup> | 133.6<sup>§</sup> | 2.55 d<sup>§</sup> |  |  |
| Minimum<sup>†</sup>: |  |  | 4960 | 2.0<sup>§</sup> | 10.2<sup>§</sup> | 76.8<sup>§</sup> | 1.40 d<sup>§</sup> |  |  |
| Maximum<sup>†</sup>: |  |  | 243 037 | 7.0<sup>§</sup> | 95.5<sup>§</sup> | 189.8<sup>§</sup> | 3.45 d<sup>§</sup> |  |  |

†: without Bismark
§: relative value per 1000 LOC

The results for the number of files, number of functions, number of code smells, and technical debt are all absolute. Therefore, it appears obvious that more code would result in higher values for all of them. To confirm this assumption about these values dependence on the project size, we did a simple linear regression for all of them with LOC as the independent variable. Table 7 contains the goodness of fit (R^2^) and the coefficients of the estimated line (slope and intercept) for each of these measurements.

**Table 7.** Linear regression results for multiple dependent variables against the lines of code (LOC)

| | $R^2$ | Slope | Intersect |
| --- | --- | --- | --- |
| **Number of files** | 0.88 | $2.8 \times 10^{-3}$ | $2.6 \times 10^1$ |
| **Number of functions** | 0.87 | $2.3 \times 10^{-2}$ | $3.3 \times 10^2$ |
| **Number of code smells** | 0.97 | $1.6 \times 10^{-1}$ | $-6.3 \times 10^2$ |
| **Technical debt** | 0.94 | $1.8 \times 10^{-3} \text{ h}$ | $2.0 \times 10^1$ |

The high values for goodness of fit (close to 1.0) confirm that both the number of code smells and technical debt have a strong linear dependency on LOC. The slopes of these two linear models suggest that with the addition of 1000 LOC about 160 new code smells are introduced on average and a time-equivalent of 1.8 days additional technical debt is to be expected. For the number of files and the number of functions, the linear model fits the measurements less precisely. Still, the values for goodness of fit confirm the assumption that more LOC means more files and more functions.

With their dependence on LOC confirmed, these absolute measurements can now be corrected for the size of the code. The distribution of their relative values (per 1000 LOC) has been plotted as boxplots that can be seen in Fig. 1. Interestingly enough, for functions per 1000 LOC the highest (95 per 1000 LOC for *MEGAHIT*) and lowest value (10 per 1000 LOC for *STAR*) match the two projects that have the lowest and second lowest debt ratio respectively. Regarding the utilisation of functions, *MEGAHIT* is marked as an outlier. It has more than double the ratio of functions to LOC than the next data point (*Bowtie* with 45 functions per 1000 LOC).

**Fig 1.**
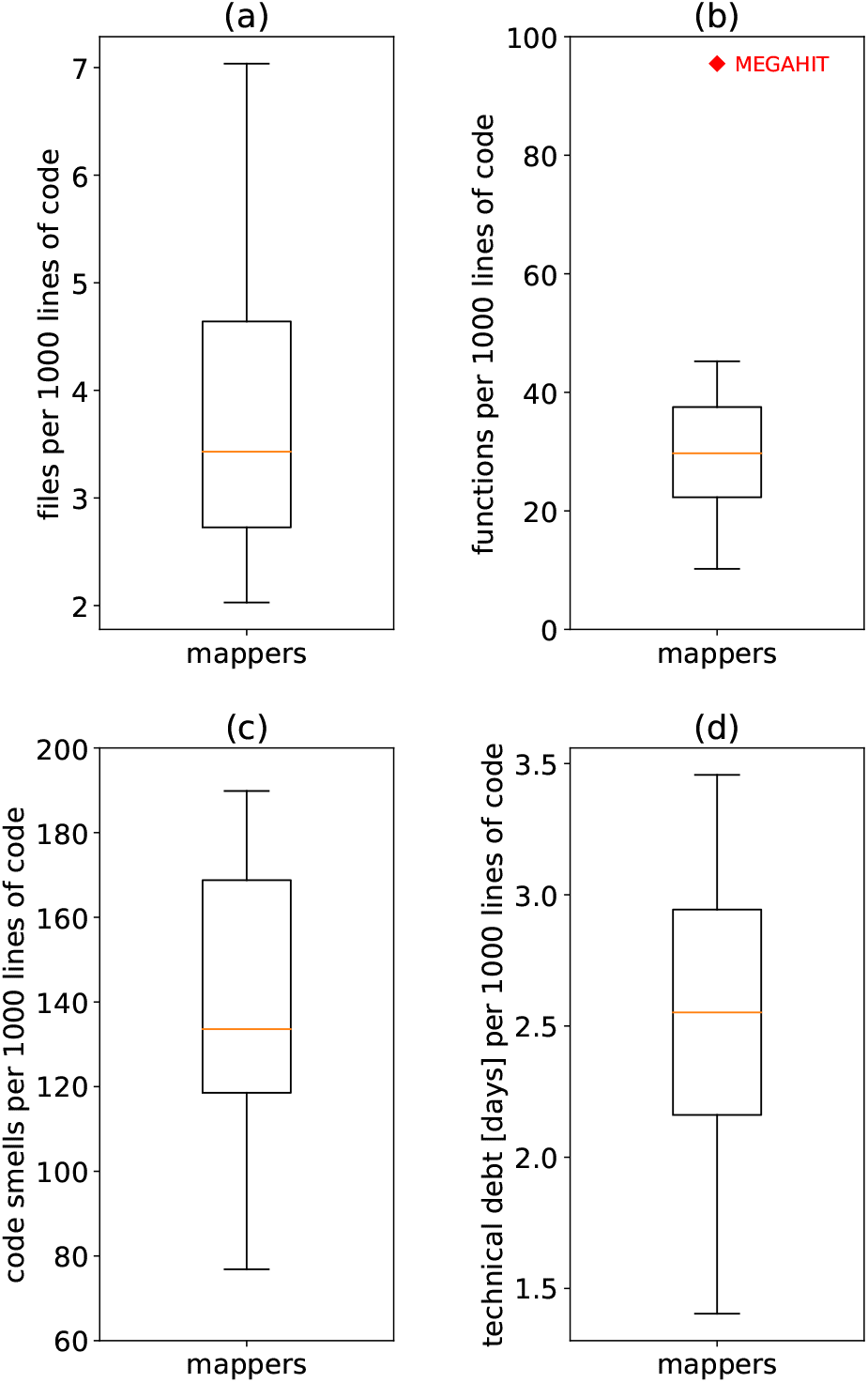
Boxplots of files, functions, code smells and technical debt (all per 1000 LOC) for the analysed group of mappers. The two upper boxplots are concerned with the structuring of the project into files (a) and functions (b). They show the medians which are about 3.4 files and roughly 30 functions per 1000 LOC. The range for files per 1000 LOC goes from 2 (*HISAT2*) to 7 (*STAR*). The lower boxplots show the number of code smells (c) and technical debt (d). The extremes in both match the debt ratio and ordering displayed in Table 6; per 1000 LOC *MEGAHIT* has the least code smells and lowest technical debt while *Bowtie* has the highest values for both.

This shows that a high level of decomposition into functions can be an indicator for a good modular code with appropriate separation of concerns. It also indicates that the average length of functions alone is a bad indicator for code maintainability. Even though Martin [61] recommends short functions that “should hardly ever be 20 lines long” and should only do one thing but do that thing well, there is a lot of discussion about this (*e.g*. [62]) and little actual data. Hence, we decided not to further interpret the information on file size and function size.

To further visualises the amount of code smells and their severity in the analysed repositories, we created a scatter plot that can be seen in Fig. 2. The graph confirms the high goodness of fit from the linear regression as all mappers are grouped closely around the line. The three most maintainable mappers, namely *MEGAHIT*, *STAR* and *TopHat*, stand out through their purple color on the technical debt colour scale. Especially for *TopHat* and *STAR* this cannot be explained through a low number of total smells but rather indicates a lower average severity of code smells; *Bowtie2* and *HISAT2* respectively have slightly less code smells per LOC and still higher technical debt per LOC.

**Fig 2.**
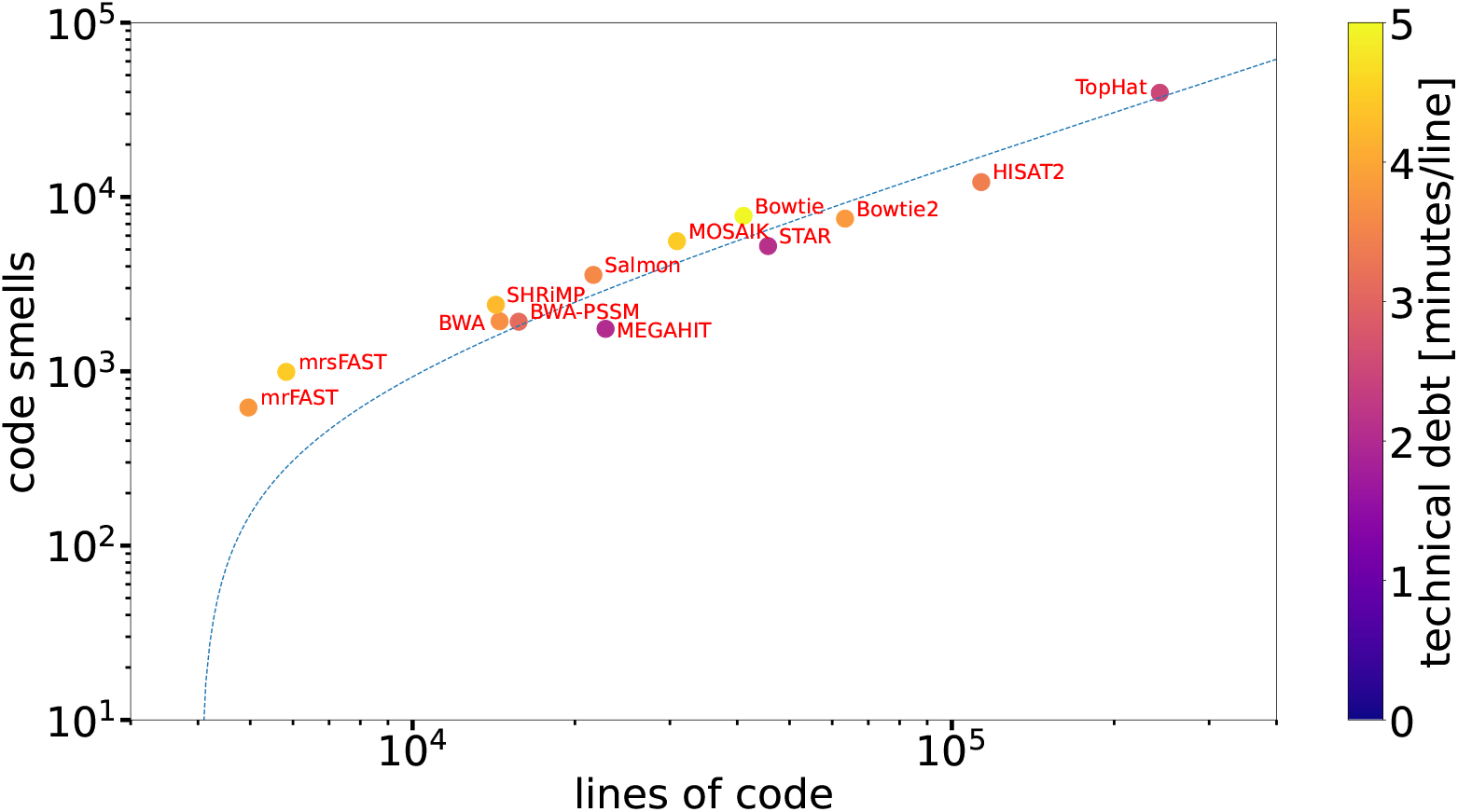
Scatterplot showing the code smells and technical debt per line of code for the analysed repositories. The color gradient is superimposed on the scatter to signal the relative technical debt of each repository. The plot shows the lines of code for each repository as an independent variable on the x-axis, while the dependent code smells are plotted on the y-axis. Since the lines of code span 2 orders of magnitude (4.7 × 10^3^ to 2.4 × 10^5^) and the number of code smells even spans 3 orders of magnitude (1.3 × 10^1^ to 4.0 × 10^4^), both variables are displayed on a logarithmic scale. Code smells are of varying severity resulting in differences in the average technical debt per code smell, therefore the technical debt per line of code contains interesting additional information. Besides the data points for the various projects, a dashed blue line is used to show the result of the linear regression for code smells against lines of code (Table 7 (*i.e*. technical debt).

Overall, the results document a very consistent degree of debt ratio (between 2.3% and 5.5%). All but 3 mappers (i.e. *mrsFAST*, *MOSAIK*, and *Bowtie*) are rated with an A, the best possible grade for their code-level maintainability. With the absolute dimensions of technical debt in these projects, this factor of 2.4 between the best (*i.e. MEGAHIT*) and worst (*i.e. Bowtie*) scoring project is quite considerable.

RQ2.1: We made the following observations about the source code maintainability of open source sequencing analysis mappers:

- Most mappers score the best possible category in the SonarSource maintainability rating
- The technical debt of mappers is independent of the project size (LOC)
- Despite low overall debt ratios, the factor of 2.4 between lowest and highest ratio is significant for absolute technical debt
- File size and function size are by themselves inadequate indicators for project maintainability

#### Process Quality

Since our observations about the process quality are entirely based on publicly available artefacts of the development process, this analysis has to be limited to tools for which such artefacts are available. This means that tools that provide their source code but do not have a publicly available version control system (i.e., an accessible git history) could not be analysed for process quality. This concerns older versions of the tools from Table 3 which were only hosted on SourceForge before moving to GitHub. The results of this analysis are summarised in Table 8. It shows the observed values for the different process quality factors described in the Development activity Section.

**Table 8.**
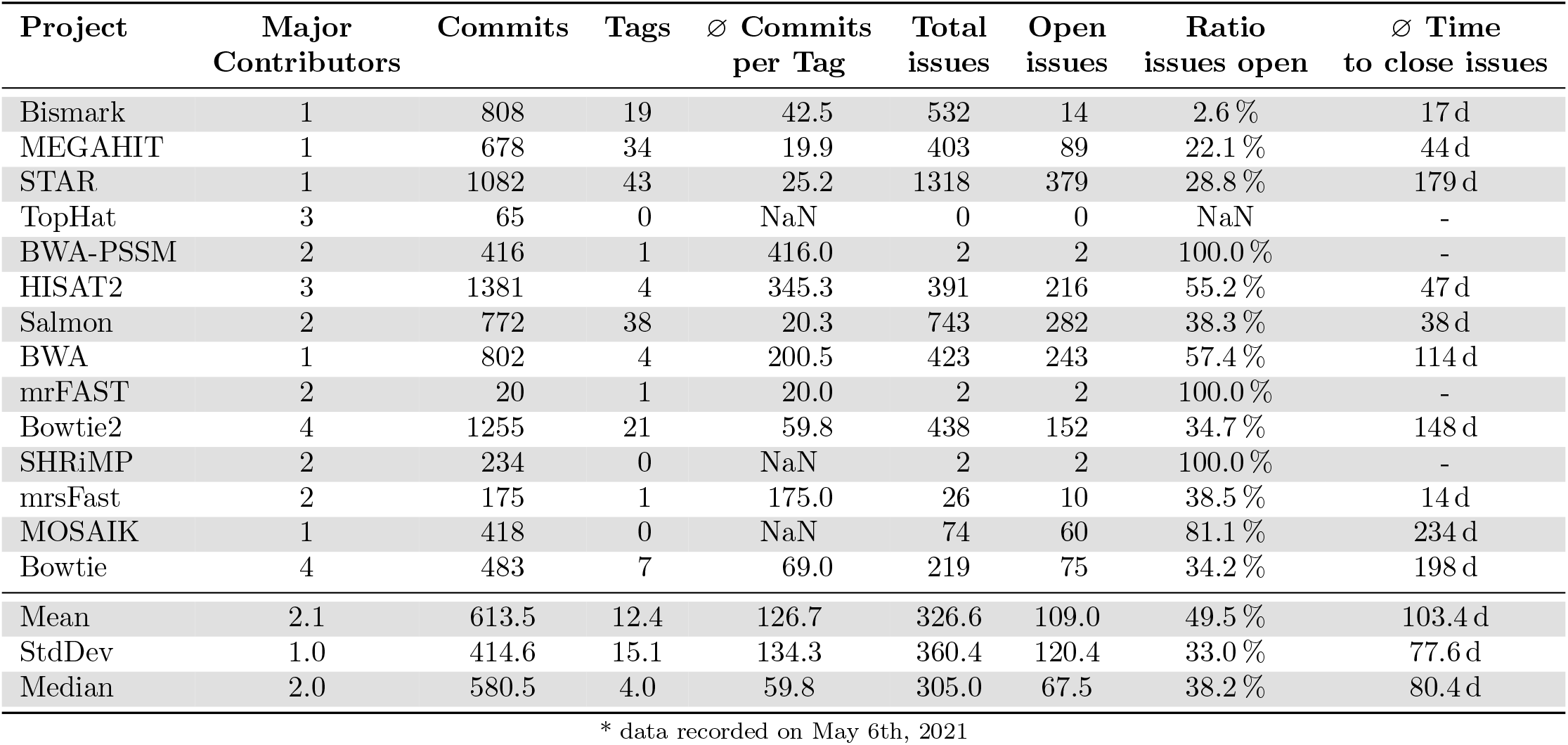
Overview of the Collected Process Quality Data.* The order of entries follows that of the maintainability data; so the technical debt ratio is ascending within this table as well.

An important result regarding the process quality is the ratio of contribution per author to the various repositories (Fig. 3). The contributors are anonymised with a capital letter and an ordinal number (e.g., A1, A2, T1).

**Fig 3.**
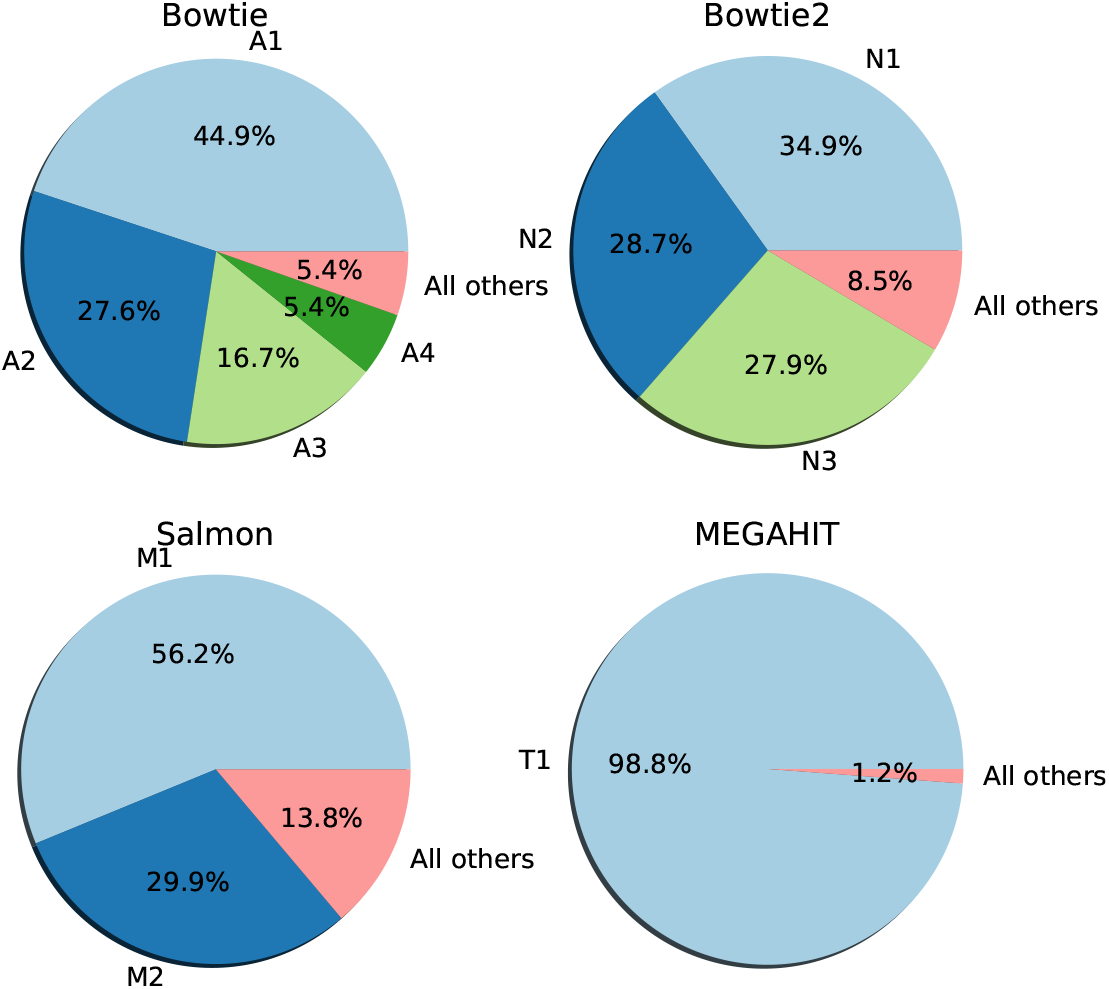
Distribution of commits to contributors for example repositories with different contribution patterns. All repositories have less than 5 main contributors.

The upper row shows examples of 2 projects (*Bowtie* and *Bowtie2*) with 4 different contributors having each contributed at least 5.0% of the total commits, and one person contributing most of the commits after 2017. The lower left plot for *Salmon* has only 2 frequent contributors. One of them is responsible for almost 60 % of the total commits, the other made over 30 % of the commits. The lower right plot for *MEGAHIT* shows another typical case where a single major contributor is responsible for the magnitude of the commits. All of the displayed examples have only a small fraction of commits made by minor contributors. The exact frequency of these four example contribution patterns is recorded in Fig. 4.

**Fig 4.**
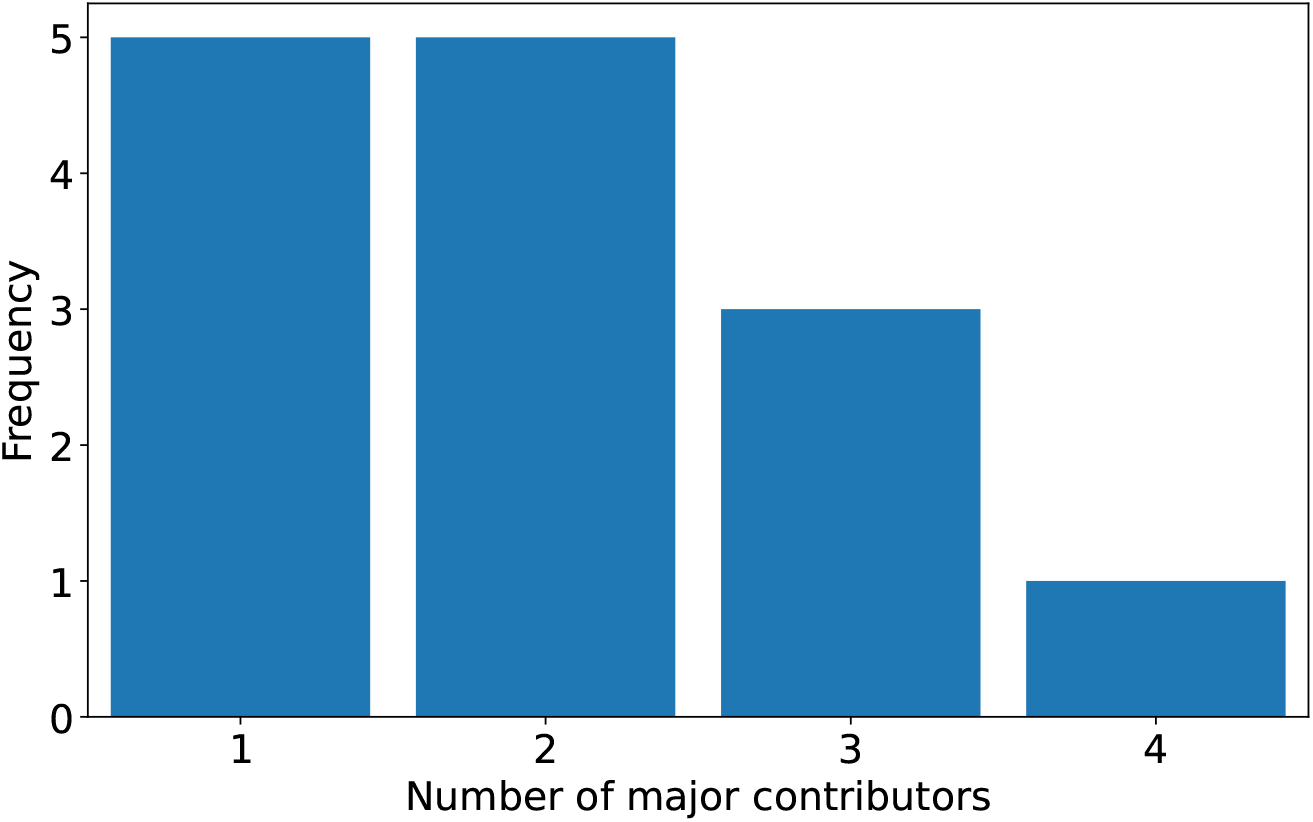
Histogram of the number of major contributors in the analysed projects (n=14).

Of the observed repositories 5, like *MEGAHIT*, have only a single major contributor. Another 5 have two major contributors (where in most cases one of them is responsible for the majority of commits) similarly to the example of *S’almon* while the remaining 4 repositories have 3 or 4 major contributors (*Bowtie, Bowtie2, HISAT2, TopHat*) and never more than 4. Especially the projects with only 1 or 2 major contributors are very dependent on the commitment of those contributors to the project. Losing those contributors means essentially that there no longer is anyone to help users with their issues. This is a significant risk that should be considered when choosing any tool.

Another interesting result in Table 8 are the number of commits, number of tags and average commits per tag. Especially the number of tags has been decisive when selecting projects for RQ2. To observe the evolution between versions, projects with sufficient number of versions were necessary. Most of the projects have fewer than 8 tags that could be used for this and are therefore not suitable for RQ2. *MEGAHIT, STAR, Salmon*, and *Bowtie2* each have over 20 tags and therefore were candidates. To visualise the timely distribution of commits and releases, we grouped commits and tags for *MEGAHIT* and *STAR* by quarterly periods and plotted them in a bar chart with dual axes. The results are shown in Fig. 5.

**Fig 5.**
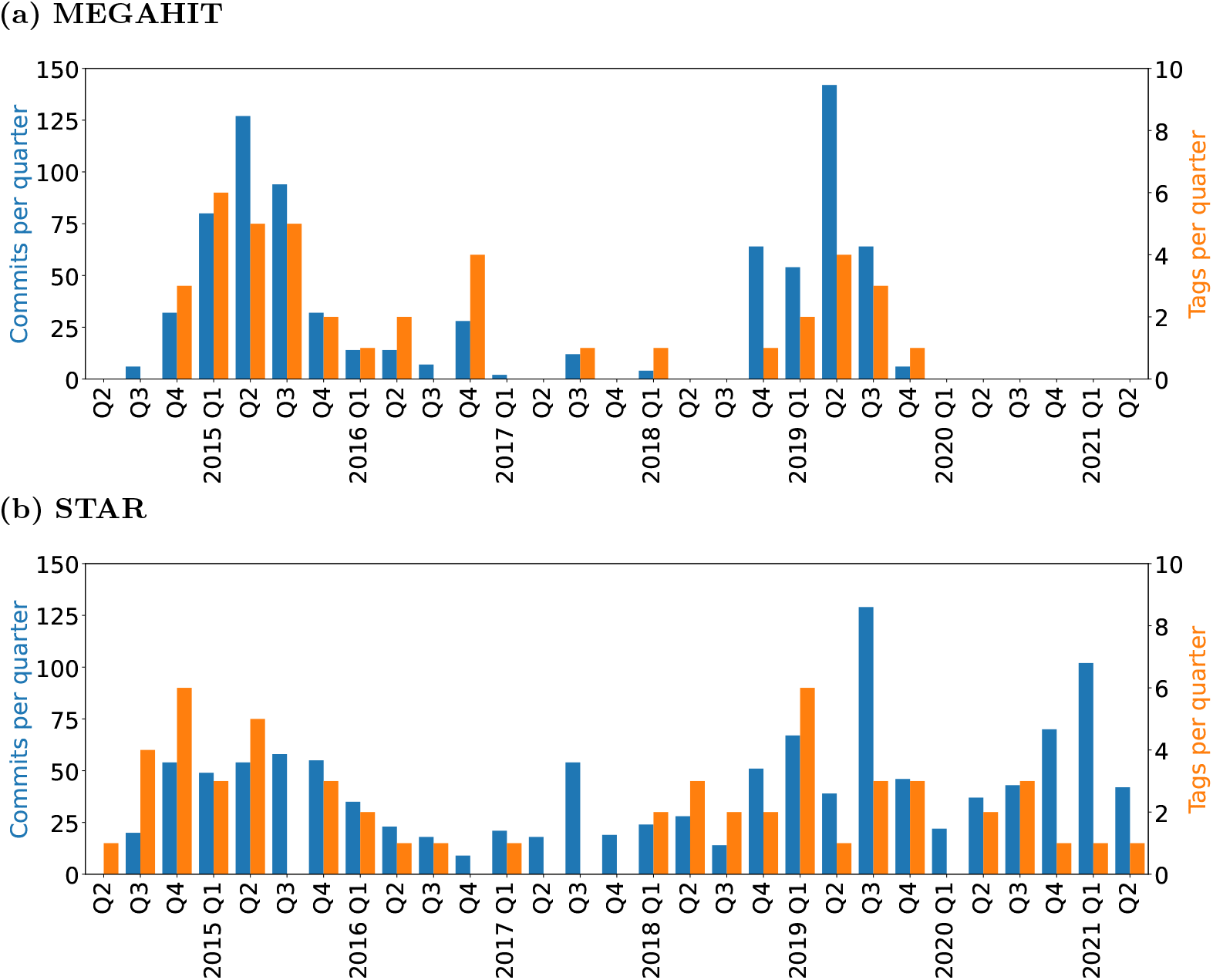
Barcharts of the quarterly commits and tags added to *MEGAHIT* (a) and *STAR* (b). The figure colour-codes two separate y-axes for commits (blue, left-side scale) and tags (orange, right-side scale). Note the factor of 15 between the two scales.

The data show more continuity in the work on *STAR* (Fig. 5b) than there is for *MEGAHIT* (Fig. 5a) over the past 7 years (both projects have been on GitHub since 2014). For *STAR* there have been commits in all quarters since Q2 2014 paired with sporadic peaks in both the coding activity and the number of tags released at those times. Most work on *MEGAHIT* in contrast was done solely during two periods of larger activity, peaking in the years 2015 and 2019. The increased number of tags generally matches the phases of increased commits in both projects.

Despite these differences in the distribution of commits and releases, *MEGAHIT* and *STAR* have a similar ratio of commits to tags with 19.9 and 25.2 commits per tag, respectively. *Salmon* (20.3 commits/tag) is in that same range and *Bowtie2* (59.8 commit/tag) has more than double the commits per tag. The numbers are still fairly similar compared to the other mappers with fewer tags, which are way beyond 100 commits per tag (e.g. *HISAT2* with 345.3) except for *Bowtie*. These projects with few tags skew the mean and standard deviation of the series to very high values. The difference between *Bowtie2* and *MEGAHIT* is not necessarily meaningful as the granularity of commits (*i.e*. changes made in a single commit) is different for different developers.

Looking at the remaining data about the issues tracked in the different projects, no clear patterns are evident. In *MEGAHIT* (22.1%) and *STAR* (28.8%) the two projects that had the highest maintainability also have low ratios of open issues. However *Bowtie* (34.2%) and *mrsFAST* (38.5%) also have low percentages of open issues and in the case of *mrsFAST* even the fastest average time to close issues once they have been opened (14 d). In this average time to close there is also a significant difference between *MEGAHIT* and *STAR*. For those issues that eventually were closed, it took an average of 1.5 months in the former project and about half a year in the latter. That constitutes a significantly longer wait for help when having issues with *STAR*. Interestingly this observation is contrary to the general consistency of development observed for both projects from Fig. 5. Despite the more frequent commits to *STAR*, issues take longer to close.

RQ2.2: The observed open source projects for sequencing analysis mappers have the following characteristics of their development activity:

- The projects are driven by few or even a single developer making major contributions
- Many of the projects are not consistently tagging updates in the code base
- The projects that consistently utilise releases average about 20 to 60 commits between consecutive releases
- Likelihood and expected time in which issues are resolved strongly vary between projects

### Evolution of Maintainability

Since software releases and tags are a main condition for the planned study of evolving maintainability, only the 4 projects with sufficient tags (# of tags ≥ 20) were candidates for this part of the study. Seeing their low technical debt ratios (see Table 6) and similarity in the observed process parameters (see Table 8), we decided to once more analyse *MEGAHIT, STAR, Bowtie2* and *Salmon* for this research question. However, we encountered build issues during the data collection of *Salmon*, thus did not include it.

The results of the static code analysis of the differently tagged versions of these 3 projects are plotted in Fig. 6. For *MEGAHIT* there is a steady increase of the LOC from the beginning to version `v1.2.0-beta` that is interrupted by 2 jumps:

1. The first jump in LOC between version `v0.2.1` and `v0.3.0-beta` is matched by a jump increasing code smells as well as minor changes in the debt ratio. This jump is caused by 127 commits being added in the course of 3 months (March 2015 to June 2015).
2. The second jump between version `v1.1.4` and `1.2.0-beta` is actually met by a drop in the number of code smells and a significantly decreased debt ratio. This second jump is caused by “heavy refactoring of the whole project” according to the main contributor. According to the changelog these changes lead to a “faster and more memory-efficient tool”. In addition to this performance boost, the overall debt ratio of the project was significantly decreased with the changes.

**Fig 6.**
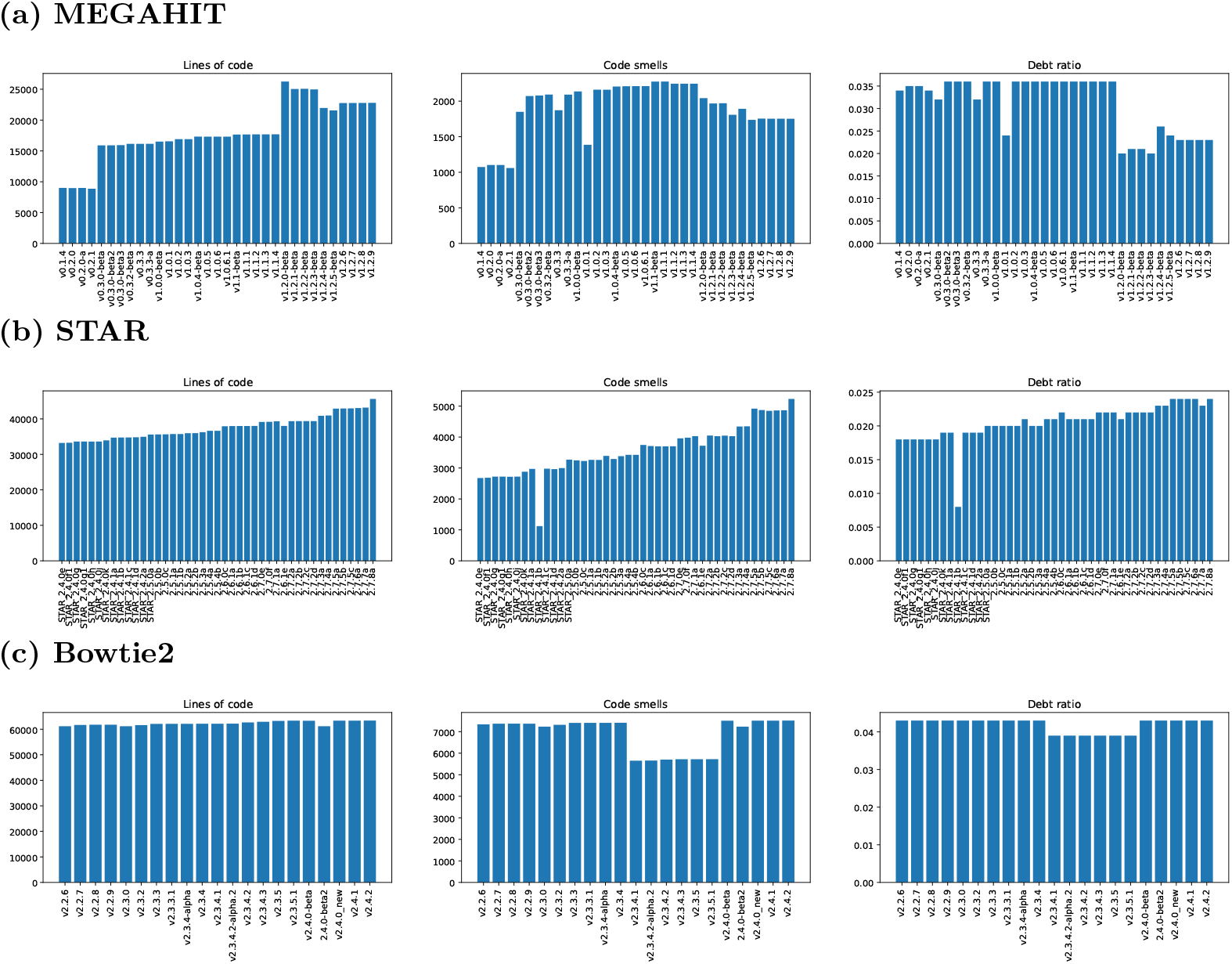
Barplots of the evolution of lines of code, code smells, and debt ratio between releases of *MEGAHIT* (a), *STAR* (b) and *Bowtie2* (c). Each of them contains three separate bar plots: The plots on the left shows the development of LOC, the central plots visualise the number of code smells, and the debt ratio of each version is visualised to the right.

In the remaining analysed releases the LOC were generally slightly reduced and so were the code smells. The debt ratio has another small jump at version `v1.2.4-beta` but was subsequently decreased again.

The plots for *STAR* do not show any special features. LOC, code smells, and debt ratio all increase steadily over the 7 years and 43 tags that have been analysed. An exception is the very low number of code smells and debt ratio in release `STAR_2.4.1b`. There have only been 4 commits changing 13 files with 58 additions and 9 deletions between `STAR_2.4.1a` and `STAR_2.4.1b`. The changes to the next version are also minor so it can be assumed that there has been an error in the analysis of `STAR_2.4.1b` that caused the drop and return in the code smells and debt ratio.

Finally, the plot for *Bowtie2* (c) almost constant values for LOC, code smells, and debt ratio. Checking the temporary drop in code smells and debt ratio for the tags `v2.3.4.1` to `v2.3.5.1` (inclusive) lead to the assumption that the change is caused by the addition (and removal in `v2.4.0-beta`) of a build flag specifying the use of `c++98`. Differences to the rules for code smells in modern `C++` probably caused the pattern.

Overall these plots show that only for *MEGAHIT* there has been an effort to consciously refactor and improve the source code. This impression is confirmed by the changelogs of the three repositories: Only *MEGAHIT* mentions refactoring, the changesin *STAR* and *Bowtie2* appear only to be concerned with fixing issues and adding functionality. A further observation from this series of analyses has been the way versions have been named in all 3 projects. The version names found on the x-ticks in Fig. 6 might sometimes follow the convention suggested by Preston-Werner [63] but they do not adhere to indicating changes according to the Major.Minor.Patch paradigm [63] most of the time. Changes in the build procedure and other breaking changes between versions have been encountered between supposed patches.

RQ3: There is no single pattern in which the maintainability of mappers evolves. We have observed a mapper that kept growing its code base and increased the technical debt in the process in *STAR*. We have observed a mapper that has not evolved much since the code is available on GitHub in *Bowtie2*. And we have observed a mapper that increased its code base at distinct times and decreased its technical debt through heavy refactoring later in the project in *MEGAHIT*.

## Discussion

### Software quality characteristics of selected tools used in HTS workflows

We found that installation and dependency management of several bioinformatics tools are poor, which should motivate the community for using conda [64] or other dependency manager software solutions. Additionally, the usability of several tools is hindered by poor error management, which can contribute to a steep learning curve limiting the access to these tools. Poor error management can also result in more time spent on a task by the end users, which hinders reaching the ultimate goal of the research. We note, that the use of Galaxy [60] enables the usage of tools for biologists who are not familiar with command line applications, thus investing in the integration of novel tools to this platform can benefit the community. Furthermore, the ease of integration of a tool into a workflow varies between tools. The challenges of integration can be partially elevated by workflow management software such as Nextflow [57], which also ensures the reproducibility of the results.

Based on these findings, we think the most effort should be focused on improving the error management of the tools by integrating solutions for the most common issues discussed on issue pages and independent forums. We also suggest improving the user experience of documentation based on the guidelines of [12]. As, ultimately, the learnability of these tools are hindered by these shortcomings, we suggest the inclusion of bioinformatics students into the maintenance process in an iterative fashion. We argue that more intuitive, easy-to-use tools with clear error messages will reduce the development time of bioinformatics workflows, thus return the investment on a global scale.

### Maintainability of selected alignment tools

In our static code analysis, we found that even though the Sonar ratings suggest a very good maintainability for most of the analysed projects, the listed technical debts are considerable. The technical debt of 32 d for *MEGAHIT* may be low compared to some of the other projects, but it still implies 96 full 8-hour working days being required to resolve that technical debt. Given that there is currently a single developer working on the project and that developer probably has further working commitments besides *MEGAHIT* (seeing the gaps in activity shown in Figure 5a), this debt is more significant than the 2.3% debt ratio reveals. Additionally, with an effective bus count of 1, the project is fully dependent on that single developer and their continued support.

The discussion on the maintainability evolution is focused on the collected data and therefore limited to versions available on GitHub. This means that only the evolution happening after initial development (*i.e*. first running versions are available) according to the software life cycle [65] can be assessed. Mostly minor new features are being developed at this point. This is especially true for *STAR* (starting from version `STAR_2.4.0e`, released 24th of October 2014) and *Bowtie2* (starting from version v2.2.6, released 30th of July 2015). Both these projects have prior development released via legacy platforms (*i.e*. Google Code and SourceForge respectively). As we are mainly interested in the maintenance phase of the software tools, this does not have a major effect on our findings.

Comparing the evolution of the technical debt between *MEGAHIT*, *STAR*, and *Bowtie2* revealed that during normal development of these projects the size of the projects (measured by LOC) only grows while the debt ratio either grows with it (in the case of *STAR*) or remains constant (in the case of *Bowtie2, MEGAHIT* before refactoring). Combining these two trends means that the absolute technical debt only grows. Brown *et al*. explain in [48] why managing technical debt is important for the long-term health of software projects:

- Technical debt is often hidden and not evident to new members of a project
- Unhandled technical debt with a project’s growth can lead to localised issues becoming widespread
- Technical debt is not necessarily growing additively; this can lead to unexpectedly reaching critical states

According to Freire *et al*. [66], there is a multitude of practices that are important to prevent technical debt. This includes project design, definition of architectural choices, project planning, creation of automated tests, and similar practices that should be obvious in modern software development. In the discussion of Table 4, we already encountered some of these as factors as they influence the maintainability of projects. Some retrospective actions could be applied to help the analysed projects improve their maintainability:

- Documenting the architectural choices that partly have been made years ago may makes it a lot easier for new developers to explore the code base
- Keeping an up-to-date list of met and open requirements tells developers and users what can be expected from a tool now and in the future
- A good test suite with high coverage makes it easier to verify the correctness of a tool after changes

Most of the analysed mappers are not employing either of these practices. A few (e.g. *Salmon*) have a test suite but the coverage is usually very limited. Especially the first two bullet points can be achieved with little effort for the expected reward. However, the one measure that our analysis shows to be very effective is refactoring. *MEGAHIT* is the mapper with the lowest debt ratio in Table 6 and this is not due to its low debt ratio throughout the development. This first place was achieved through extensive refactoring between the two versions. Between these versions the technical was reduced by 6 days while almost 9000 LOC were added. Had those 9000 lines been added without improved debt ratio, the technical debt would have been 25 days higher than it actually was with the refactoring. This shows that a dedicated refactoring effort – like the work done on *MEGAHIT* over a period of 5 month – is realistic in bioinformatics tools and can have a significant impact on the maintainability of a project. In their systematic review of research on code smells and refactoring, Lacerda *et al*. also come to the conclusion that refactoring should be the first measure when attempting to reduce technical debt [67].

### Improving the Development Process

The origin of the term “bus count” suggests that a high number of active developers should be an ambition of any open source software tool. More active developers who are familiar with the project does not only mean that a project can be continued even if one of them stops to work on it. We also expect that projects with more active developers should be faster in their response to the issues being reported. This assumption was however not confirmed by the data. Neither the ratio of open issues nor the average time to close them was found to generally be better for projects with 3 or 4 major contributors compared to those with only 1 or 2. These values turned out to be different on a case by case basis.

A further advantage of a larger number of involved contributors is that it allows for the implementation of pull requests as a mean of peer reviewing code. According to Silva *et al*. pull requests can be a means of reducing technical debt continuously throughout the development process if used in the right way [68]. The low number of major contributors combined with the additional effort required for proper code reviews, however, makes this an unrealistic solution in the given scenario. An automated approach without the human factor is recommended instead. We therefore recommend adding the following tools and steps to the development process:

- A linter to enforce strict formatting and further programming rules. Research has shown linters to be effective in reducing code smells and vulnerabilities [69]
- Proper use of semantic versioning [63] and a changelog [70]
- Projects with frequent changes and sufficient test coverage can also benefit continuous integration

In information technology *linters* are tools used to detect issues in code. This includes stylistic and semantic issues. Semantic versioning is the usage of the software version number in the following way: MAJOR.MINOR.PATCH. MAJOR increments when an incompatible API change is made, MINOR increments when a new feature is added which is backward compatible to previous changes, and PATCH increments when a bug is fixed in a backward compatible way. Continuous integration is the process of iteratively adding new code to a working code base, while making sure that no new code is causing breaking changes.

These are some easy steps that can be applied even to projects run by a single developer. They will help reduce technical debt, keep the users and co-developers updated about changes and compatibility between releases, and prevent publishing of changes that break tests or build procedures.

## Conclusion

The data collected in this research shows that bioinformatics mappers are generally at a good software quality and maintainability level. However, code quality shows a trend of degradation over time, which can be reversed with a conscious effort of refactoring. The development of the investigated software is usually driven by very few major developers, creating a strong dependency on those developers’ commitment to the projects. This not only results in varying success of handling issues in the code base, but hinders refactoring efforts too. We therefore recommend a set of practices that can easily be implemented even in projects with a single major contributor and should help to steadily and permanently improve the maintainability of open source mappers and other tools for scientific computation.

With the continuous development of scientific software, we would like to see further research into the implementation and effects of the recommended improvements. The tooling we provide makes the collection of future data on the subject very easy and we hope it can be used to assess the future development of the analysed mappers.

## Acknowledgment

MD was supported by the Swedish Foundation for Strategic Research (RIF14 · ®0081)

## Footnotes

1 https://sonarcloud.io

