## Supplementary figures and images for "Empirical study on software and process quality in bioinformatics tools"

### Supplementary Fig.1

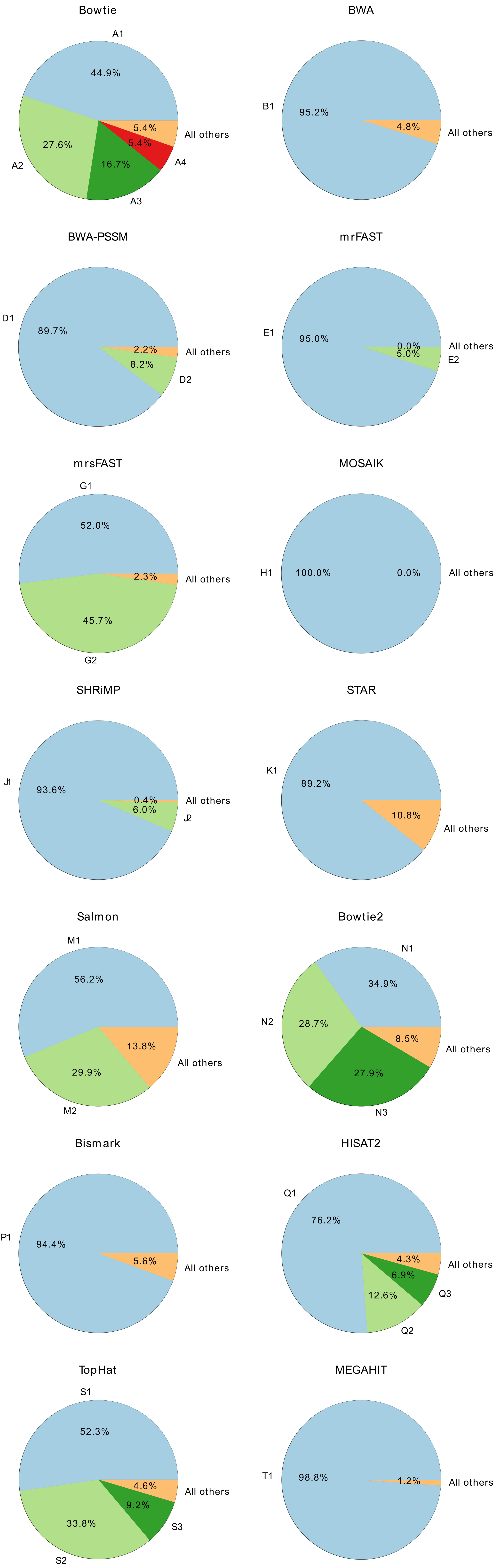
